# Towards a Generative Paradigm for Large-scale Microbiome Analysis by Generative Language Model

**DOI:** 10.1101/2025.01.15.633278

**Authors:** Haohong Zhang, Zixin Kang, Yuli Zhang, Ronghua Yang, Kang Ning

## Abstract

Microbiome analysis has traditionally relied on taxonomic abundance tables, which, while effective, often constrain the exploration of deeper contextual relationships. In this study, we present MGM 2.0, a novel framework that applies advanced natural language processing (NLP) techniques to microbiome research. By reimagining microbiome samples as sentences and microbial species as words, MGM 2.0 enabled the extraction of nuanced patterns and relationships. The model demonstrated robust predictive performance in identifying exogenous species colonization (AUROC = 0.86). Additionally, through prompt-guided microbiome data generation, MGM 2.0 produced realistic microbial profiles conditioned on disease labels. The framework further revolutionized donor selection in fecal microbiota transplantation (FMT) by framing it as a sequence-to-sequence prediction task, enabling the prediction of post-transplantation community compositions and the identification of super donors for personalized treatments (average increase in C2R = 0.52). This innovative integration of NLP and microbiome science provides a versatile toolkit for predictive modeling, data generation, and personalized medicine.

**Highlights:**

- Introduced MGM 2.0, a generative language model utilizing sentence-like representation for microbiome analysis and generation.
- Demonstrated that sentence-like representation preserves sample distinctions, enabling accurate microbial sample classification tasks, such as colonization prediction.
- Generated realistic, disease-specific microbiome profiles using a prompt-guided approach, validated by a novel “Microbiome Turing Test.”
- Applied MGM 2.0 to fecal microbiota transplantation (FMT) donor selection, accurately predicting post-transplant community compositions and identifying potential “super donors” for personalized treatment strategies.

## Introduction

Microbiome analysis has been instrumental in elucidating the composition, functions, and interactions of microbial communities across diverse environments [1, 2]. Traditionally, taxonomic profiling abundance tables have served as the primary data representation, summarizing the relative proportions of microbial taxa within a sample [3, 4]. While this approach effectively captures quantitative relationships, it often neglects the intricate contextual dependencies between microbial taxa and their communities, which restricts the potential of microbiome data to drive predictive and functional insights due to its sparsity, overdispersion and high-dimensionality [5-7]. Recently, several methods implemented phylogenetic information as a restriction to enhance microbial analysis [8-10]. However, despite these methods improve the performance of prediction task, the high-level task like data generati did not benefit from the integration of information from such approach.

Recent advancements in natural language processing (NLP) offer a promising avenue to overcome these limitations [11]. By treating tabular data through the lens of sentence-like representations, analogous to text in NLP, we can leverage sophisticated computational frameworks that excel at modeling context and relationships [12-14]. In our previous study, we introduced the Microbial General Model (MGM), a microbiome foundation model that transforms species abundance tables into sentence-like representations, treating microbiome samples as “sentences” and microbial taxa as “words” [15]. MGM was pretrained on a large-scale corpus of 260,000 microbiome samples, enabling it to encode rich contextual information and capture complex inter-species interactions. This foundational framework provided a significant leap in microbiome research by leveraging state-of-the-art natural language processing (NLP) techniques such as self-supervised learning and attention mechanisms.

Building on this foundation, we developed MGM 2.0, which enhances the generative capabilities of microbiome modeling. This upgraded framework introduces two transformative innovations: (1) generating realistic microbiome abundance profiles from prompts, facilitating controlled in silico experiments and simulation, and (2) predicting post-transplant microbiome compositions by modeling donor and recipient communities in fecal microbiota transplantation (FMT). By incorporating advanced NLP techniques, including prompt-based learning and sequence-to-sequence generation, MGM 2.0 transcends static data representations to enable dynamic, context-aware applications. These advancements hold immense potential for personalized medicine, ecological forecasting, and synthetic biology, highlighting the versatility and transformative impact of NLP-driven approaches in microbiome research.

## Results

### NLP techniques enabled deep understanding of microbial community

MGM 2.0 leverages natural language processing (NLP) techniques to redefine microbial community analysis. By employing sentence-like data representations, it transforms microbial abundance data into discrete input formats through genus-level normalization, tokenization, and the ranking of taxa by relative abundance (**Fig. 1a**). Traditional machine learning algorithms have demonstrated strong performance in modeling microbial abundance tables, including linear methods such as LASSO [16], tree-based methods like Random Forests (RF) [17], and deep learning approaches like fully connected neural networks [18]. While these models primarily rely on supervised learning to capture inter-sample distinctions in labeled datasets [19, 20] , MGM 2.0 employs self-supervised strategies inspired by NLP to focus on individual samples and the intricate structures within microbial communities (**Fig. 1b**) [21, 22].

**Figure 1.**
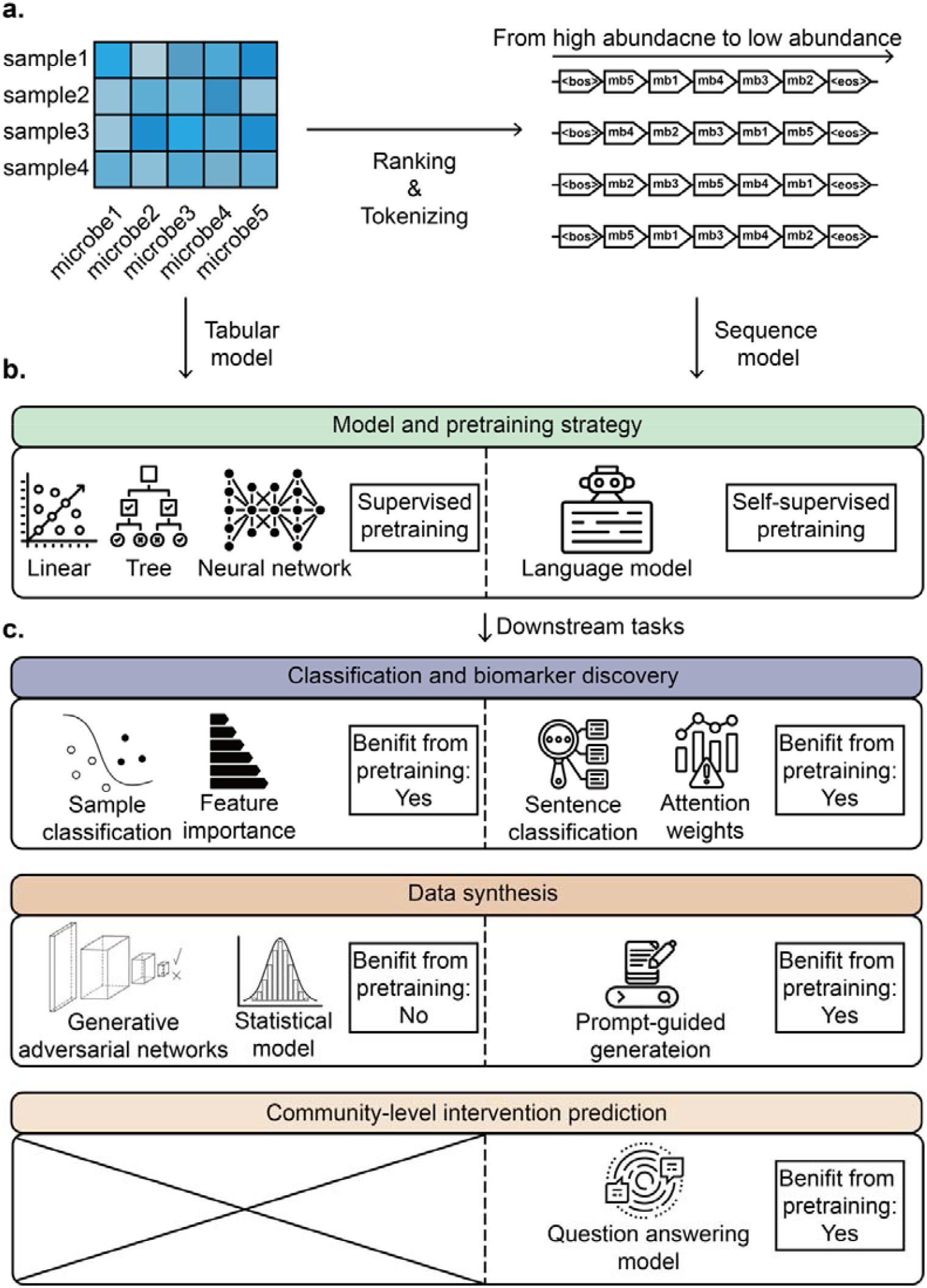
Comparation between the taxonomic profiling abundance table and sentence-like representation of microbial community. **a.** Tabular-to-sequence transformation process. **b**. Common methods and pretraining strategy to handle the tabular or sequential data. **c**. Specific methods on common microbiome problems and whether it benefits from pretraining.

The integration of NLP methodologies enables common microbiome tasks, such as sample classification and biomarker discovery, to align with analogous tasks in NLP, including [23] and attention weight analysis [24]. Since these tasks focus on inter-sample differences, both supervised and self-supervised pretraining can enhance model performance. For generative tasks, MGM 2.0 demonstrates a notable advantage over traditional methods such as statistical models and Generative Adversarial Networks (GANs) [25]. These conventional approaches are constrained by their focus on the compositional characteristics of individual samples and derive limited benefits from supervised pretraining. In contrast, MGM 2.0 utilizes prompt-guided generation, leveraging pretrained models that capture microbial community structures without biases introduced by labels [21]. This capability is particularly powerful for generating realistic microbiome profiles conditioned on specific prompts, enabling in silico experimentation and hypothesis testing. Additionally, NLP techniques facilitate the prediction of community-level interventions through sequence-to-sequence modeling, addressing challenges that traditional abundance table-based approaches struggle to overcome. By incorporating NLP-driven methodologies, MGM 2.0 sets a new standard for microbial community modeling, offering transformative capabilities for predictive and generative microbiome research (**Fig. 1c, Supplementary Table 1**).

**Table 1.** Microbiome Turing Test of Different Methods. This table summarizes the performance metrics of three different methods: MGM, MB-GAN, and Random, along with a baseline for comparison. Each metric is reported with its mean value and 95% confidence interval where applicable.

| Metrics | True | MGM | MB-GAN | Random |
| --- | --- | --- | --- | --- |
| Phase 1 |  |  |  |  |
| Cosine Similarity | - | **0.52±0.069** | 0.046±0.0077 | 0.05±0.0084 |
| MAE | - | 0.00013±1e-05 | 0.00011±8.3e-07 | 0.00011±6e-07 |
| Alpha diversity | 2.8±0.33 | **3.4±0.3** | 1.5±0.055 | 1.6±0.026 |
| Sparsity | 1.9±0.23 | **2.4±0.24** | 1.1±0.038 | 1.1±0.018 |
| Spearman | - | **0.51±0.05** | 0.0018±0.00073 | 0.0015±<br>0.00063 |
| Phase 2 |  |  |  |  |
| ROC-AUC | - | **0.95** | 0.49 | 0.5 |
| Biomarkers Overlap<br>(Top 500) | 500 | **440** | 287 | 283 |
| Network Density | 0.18 | 0.30 | 0.10 | 0.10 |
| Network average<br>degree | 281.16 | **351.88** | 959.98 | 959.98 |
| Network modularity | 0.22 | **0.16** | 0.13 | 0.13 |

| Metrics | True | MGM | MB-GAN | Random |
| --- | --- | --- | --- | --- |
| Phase 1 |  |  |  |  |
| Cosine Similarity | - | **0.52±0.069** | 0.046±0.0077 | 0.05±0.0084 |
| MAE | - | 0.00013±1e-05 | 0.00011±8.3e-07 | 0.00011±6e-07 |
| Alpha diversity | 2.8±0.33 | **3.4±0.3** | 1.5±0.055 | 1.6±0.026 |
| Sparsity | 1.9±0.23 | **2.4±0.24** | 1.1±0.038 | 1.1±0.018 |
| Spearman | - | **0.51±0.05** | 0.0018±0.00073 | 0.0015±<br>0.00063 |
| Phase 2 |  |  |  |  |
| ROC-AUC | - | **0.95** | 0.49 | 0.5 |
| Biomarkers Overlap<br>(Top 500) | 500 | **440** | 287 | 283 |
| Network Density | 0.18 | 0.30 | 0.10 | 0.10 |
| Network average<br>degree | 281.16 | **351.88** | 959.98 | 959.98 |
| Network modularity | 0.22 | **0.16** | 0.13 | 0.13 |

### Sentence-like representation effectively captures microbial variations across samples

To assess whether our sentence-like representation method retains the differential information between samples, we conducted an evaluation using a dataset from an exogenous species colonization experiment [26]. This dataset comprised 24 individuals, each subjected to antibiotic intervention and subsequently exposed to *Enterococcus faecium* (*E. faecium*) as the exogenous species. We aimed to evaluate whether sentence-like representation could effectively capture differential information across microbial communities by comparing the performance of two models: a RF model trained on the original species abundance table, and a MGM sentence classification model, which was applied to both classification and regression tasks.

In the classification task, where the goal was to predict whether *E. faecium* would successfully colonize a given sample, the MGM model achieved comparable performance to the RF model, with an accuracy of 0.86 versus 0.83, respectively, based on five-fold cross-validation (**Fig 2a, b**). This result suggests that the sentence-like representation employed by the MGM model effectively preserves the essential differential information between samples, enabling it to capture variations in microbial communities despite the transformation from raw abundance data to sequence-based representations.

**Figure 2.**
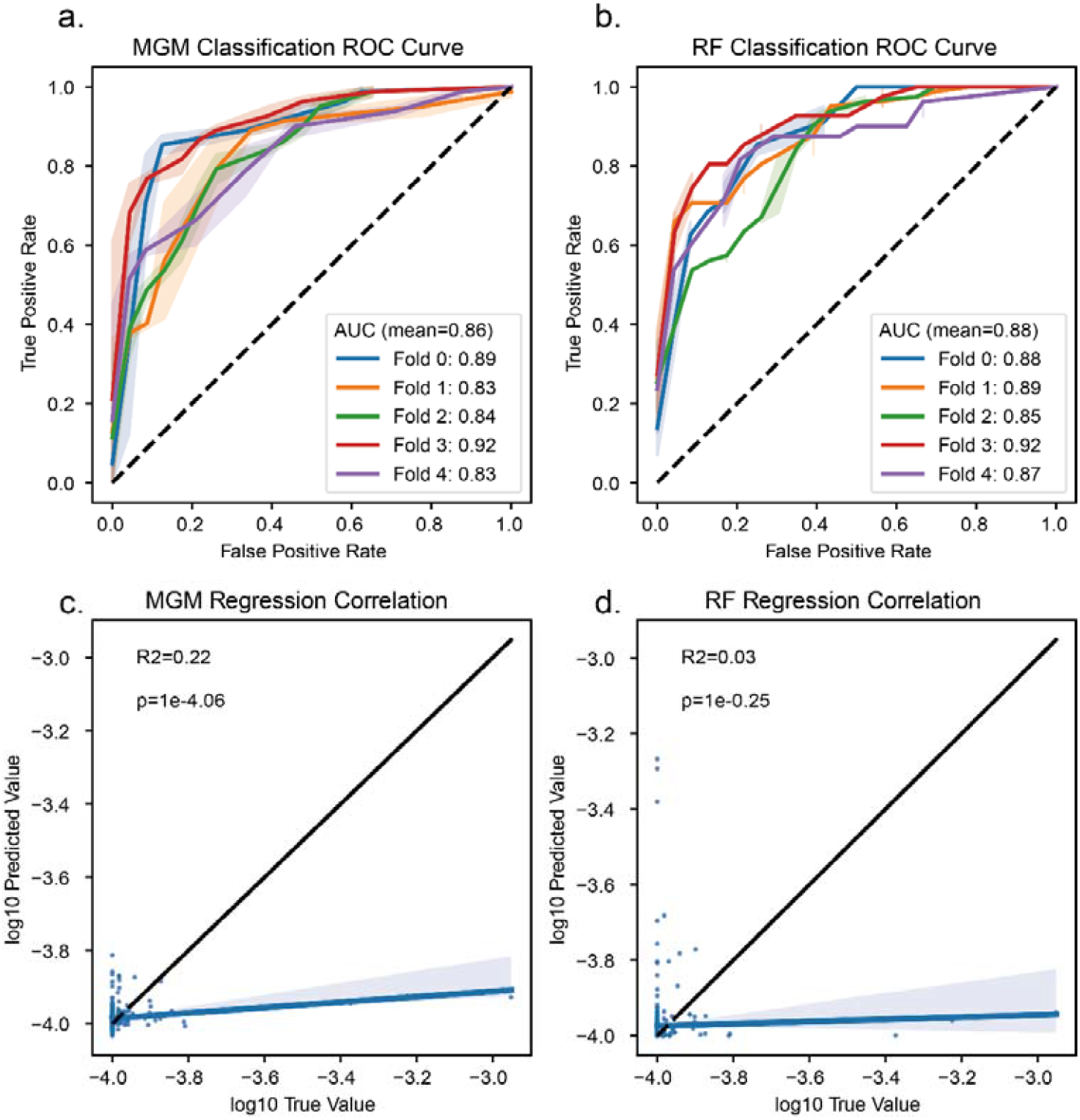
The predicted colonization outcomes of *E. faecium*. **a**. ROC curve of MGM sentence classification models in binary classification (permissive vs. resistance) of the colonization outcomes of *E. faecium*. **b**. ROC curve of random forest models in binary classification (permissive vs. resistance) of the colonization outcomes of *E. faecium*. **c**. Correlation of MGM sentence classification models in regression of the colonization outcomes of *E. faecium*. **d**. Correlation of RF models in regression of the colonization outcomes of *E. faecium*.

The regression task, aimed at predicting the post-intervention abundance of *E. faecium*, produced even more striking results. Here, the MGM model significantly outperformed the RF model, achieving an R^2^ value of 0.22 compared to 0.03 for RF (**Fig 2c, d**). This outcome indicates that, despite the sentence-like transformation losing some of the original abundance information, the MGM model was still able to capture sufficient underlying structure to predict *E. faecium* abundance post-intervention.

These findings demonstrate that sentence-like representation holds promise as an effective approach for colonization prediction tasks. The MGM model successfully retained key differential information, allowing it to perform robustly even when direct abundance values were not preserved.

### Prompt-guided microbiome generation and evaluation via a Microbiome Turing Test

To showcase the generative capabilities of MGM 2.0, we developed a prompt-guided pipeline to synthesize realistic microbiome abundance profiles. Given that our sentence-like encoding omits original relative abundance information, we implemented a reconstructor network to convert generated rank sequences back into abundance tables.

The generation process involved augmenting the model’s vocabulary with label tokens appended after the beginning-of-sequence token (’<bos>’) for each sample, followed by fine-tuning using next-token prediction. For example, the sentence-like representation of a sample labeled ‘CRC’ included a ‘CRC’ token following the ‘<bos>’ token. During generation, each label token served as a prompt, generating microbiome sequences with a length consistent with the original samples. These sequences were then transformed into abundance tables using the reconstructor (**Fig. 3a, Methods**).

**Figure 3.**
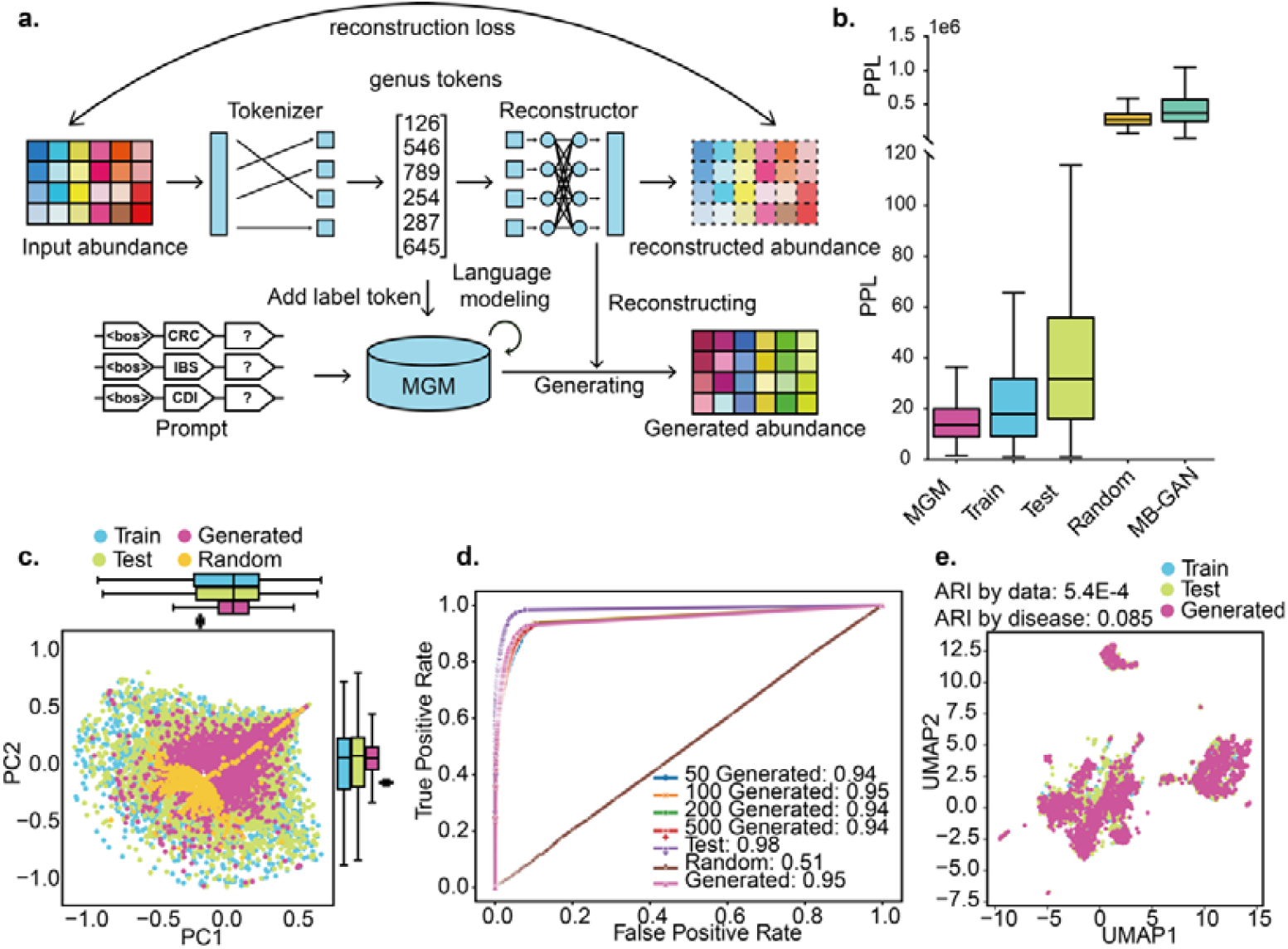
Evaluation of the prompt-guided MGM generative model. **a**. MGM generative model pipeline. **b**. Perplexity of different datasets. Perplexity in language models measures the degree to which the model considers a sentence to be realistic. **c**. Beta diversity analysis of the reconstructed model, based on Bray-Curtis distance. **d**. ROC-AUC curves of the classifier for gradient-generated data. **e**. UMAP dimensionality reduction plots of embeddings for three data types (train, test, generated). The ARI indices represent clustering based on data type and disease type, with clustering labels obtained through a Gaussian Mixture Model.

We utilized 6,004 gut microbiota samples representing 17 diseases from the GMrepo database [27] to train and evaluate our generative model. The dataset was split into equal training and testing sets using hierarchical sampling. Initial evaluation focused on perplexity (PPL), a measure of how well the model predicts a sequence. Generated samples exhibited the lowest perplexity compared to training, external test, and randomly generated samples (**Fig. 3b**), indicating high realism. Beta diversity analysis, based on Bray-Curtis distance, further confirmed that the distribution of generated samples closely mirrored that of real data (**Fig. 3c**). To assess the biological relevance of the generated data, we trained a sentence classification model for the 17 diseases. This classifier achieved a ROC-AUC of 0.98 on the test data and 0.95 on the generated data, demonstrating that the generated samples retained disease-specific information. We also generated varying numbers of samples per disease (50, 100, 200, and 500) and found consistent performance of the disease classifier across these gradient-generated datasets (**Fig. 3d**), confirming the robustness of our generative model. UMAP dimensionality reduction of embeddings further revealed that generated samples clustered more strongly by disease type (ARI=0.085) than by data type (train, test, generated; ARI=5.4E-4), highlighting the model’s ability to capture disease-specific characteristics (**Fig. 3e**).

To comprehensively evaluate the generative capabilities of MGM 2.0, we developed a novel assessment framework: the Microbiome Turing Test. This framework consists of two phases: evaluating the statistical properties of the generated data and assessing its biological significance.

In the first phase, we examined the statistical properties of the generated microbiome data. Alpha diversity was calculated using Shannon’s index to assess species richness and evenness within samples. The Mean Absolute Error (MAE) between the relative abundances of real and generated samples quantified the average deviation, while Cosine Similarity measured the resemblance in community composition by evaluating the cosine of the angle between abundance vectors. Sparsity was analyzed by comparing the proportion of zero entries in abundance matrices, reflecting the similarity of sparsity patterns in real and generated data. Finally, Spearman Correlation coefficients between taxa abundances were computed to capture monotonic relationships within microbial communities.

The second phase focused on evaluating the biological relevance of the generated data. A random forest classifier trained on real data achieved high ROC-AUC scores when applied to generated data, confirming the retention of disease-specific microbial patterns. Additionally, a classifier trained on the generated data exhibited significant overlap in the top 500 important microbes with classifiers trained on real data, further validating the preservation of key biomarkers. Relative network analysis was conducted to explore functional groupings and interactions within the microbiome. Co-occurrence networks were constructed, with nodes representing microbial taxa and edges denoting significant associations. Metrics such as network density (ratio of observed to possible edges), average degree (typical connectivity per node), and modularity (extent of distinct community structures) were used to compare the generated and real data.

To benchmark MGM 2.0, we compared its performance against MB-GAN, a GAN-based microbiome data generation approach [28]. Across multiple metrics of the Microbiome Turing Test, MGM 2.0 demonstrated superior performance (**Table 1**). Unlike MB-GAN, which requires retraining for each disease, MGM 2.0 leverages a prompt-guided approach, enabling efficient and flexible data generation conditioned on specific prompts. This advantage significantly enhances its utility for targeted microbiome research and in silico experimentation.

### Question answering strategy enabled predicting donor fitness in fecal microbiota transplantation

To investigate the MGM 2.0’s generative capabilities for microbiome applications, we addressed donor selection in fecal microbiota transplantation (FMT) by predicting community-level perturbations. We framed FMT as a question-answering task, analogous to those in natural language processing. Specifically, recipient and donor sequence representations were concatenated to form the input query, and the model generated a predicted post-transplantation community composition using a sequence-to-sequence (seq2seq) approach. For example, given an FMT triad of recipient 1, donor 1, and post-transplantation sample 1, the input query consisted of the concatenated sentence-like representations of recipient 1 and donor 1, separated by a ‘sep’ (separate) token, with the sentence-like representation of post-transplantation sample 1 serving as the target sequence (**Fig. 4a, Supplementary Table 1**). We fine-tuned the model on a dataset of 228 FMT experimental groups [29] and validated it using two external datasets: Khanna et al. (15 FMT pairs from inflammatory bowel disease (IBD) patients) [30], and Goyal et al. (38 FMT pairs from *Clostridioides difficile* infection (CDI) patients) [31]. Performance was evaluated using the ROUGE-1 metric, achieving consistent scores exceeding 0.6 across training, validation, and testing datasets, with particularly strong performance observed on the Khanna et al. dataset (**Fig. 4b**).

**Figure 4.**
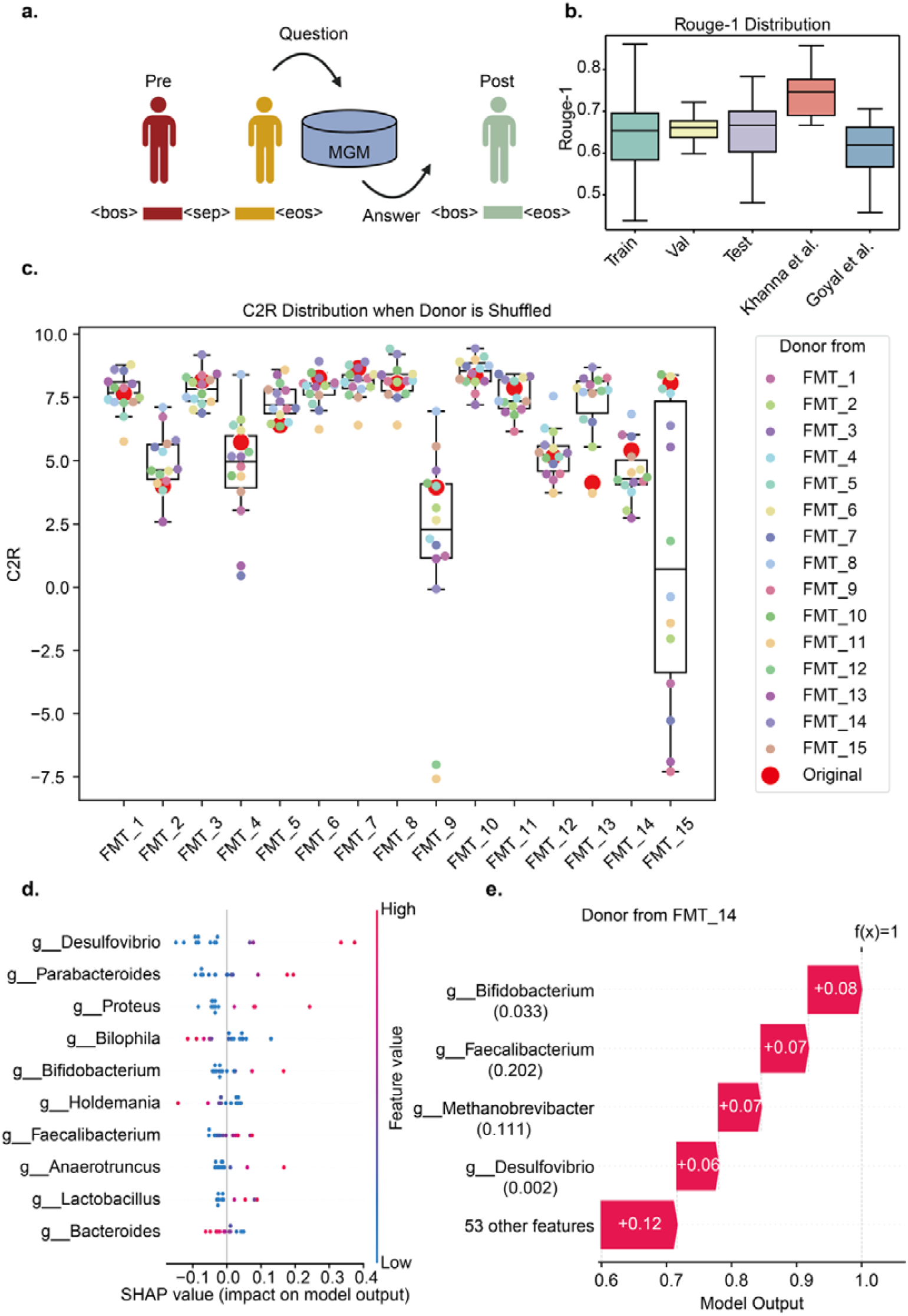
Pipeline and evaluation of the community-level intervention prediction. **a**. The FMT experiment could be abstracted as a question-answering task. **b**. Rouge-1 evaluation on both inner and external dataset. **c**. C2R distribution of in silico reassignment of donors to different recipients. The large red points indicate the original donor-recipient pairs. **d**. SHAP summary plot highlighting genus-level contributors to C2R values. **e**. Waterfall plot explaining the SHAP values for the donor from FMT14.

To assess our model’s utility for donor selection, we performed an additional validation using the Khanna et al. dataset. We systematically reassigned donors to different recipients within this dataset to generate alternative transplantation scenarios. The model predicted the resulting community composition for each new recipient-donor pairing. Community engraftment success was quantified using the Community-to-Recipient (C2R) metric. While the originally paired donors exhibited high C2R values, our analysis revealed that they were not always the optimal choice. Notably, the donor from FMT14 consistently yielded the highest C2R values across multiple recipient groups, suggesting potential as a “super donor” (**Fig. 4c, Supplementary Fig. 1**, Average increased C2R = 0.52). Conversely, the donor from FMT11 consistently resulted in the lowest C2R values when transplanted to other recipients, although it performed well in the original pairing (**Fig. 4c, Supplementary Fig 1**, Average increased C2R = -2.20).

To investigate the factors contributing to donor effectiveness, we developed an LDA model to classify donors as having positive or negative impacts on C2R outcomes and employed SHapley Additive exPlanations (SHAP) [32] analysis to identify genus-level drivers influencing FMT success (**Fig. 4d, e; Supplementary Fig. 2a-e**). This analysis revealed that donors with higher relative abundances of genera such as *Desulfovibrio* and *Bifidobacterium* were strongly associated with elevated C2R values, indicating enhanced community engraftment potential. Conversely, genera such as *Bilophila* and *Holdmania* were negatively associated with C2R values, suggesting they may hinder successful transplantation (**Fig. 4d**). Notably, previous studies have suggested that utilizing donor feces rich in *Bifidobacterium* can stimulate the recipient’s microbiota to recover from decreased diversity to levels comparable to the donor’s microbiota [33].

A detailed examination of the donor from FMT14, visualized using a waterfall plot, showed that *Bifidobacterium* and *Faecalibacterium* had a significant positive impact on the model output (+0.08 and +0.07, respectively) and were present at relative abundances of 0.033 and 0.202, respectively (**Fig. 4e**). These findings suggest that the enrichment of these genera may be critical to this donor as a “super donor.” These results reinforce the importance of specific microbial taxa in FMT success and underscore the potential of MGM 2.0 to provide interpretable and actionable insights into microbiome-based therapeutic strategies.

By leveraging a question-answering framework inspired by natural language processing, MGM 2.0 offers a scalable, accurate, and interpretable method for advancing microbiome-based therapeutic interventions. These findings highlight its promise in optimizing donor selection and improving outcomes in FMT.

## Discussion

This study introduces MGM 2.0, a generative language model for microbiome analysis and generation. By transforming microbiome data into sentence-like representations, MGM 2.0 enables the application of advanced NLP techniques, facilitating deeper contextual understanding and enhanced predictive capabilities.

Our findings demonstrate the efficacy of MGM 2.0 across diverse tasks. The sentence-like representation preserves crucial sample-specific information, achieving classification accuracy comparable to traditional methods while significantly enhancing regression performance. This showcases the model’s ability to capture and utilize nuanced patterns and dependencies within microbial communities, underscoring the value of adapting NLP methodologies to microbiome research.

A key innovation of MGM 2.0 lies in its robust generative capabilities for microbiome data generation. Through prompt-guided generation, the model produces realistic, biologically meaningful profiles conditioned on specific contexts, such as disease states. Fine-tuning a pre-trained model, combined with a reconstructor network, enabled us to generate profiles that closely mimic real-world distributions. The Microbiome Turing Test—a comprehensive evaluation framework incorporating biological significance and data distribution metrics—validated the superiority of MGM 2.0 over methods like MB-GAN. This capability is particularly valuable for generating synthetic datasets, which can augment limited real-world data, facilitate machine learning model development, and explore hypothetical microbiome states. Moreover, MGM 2.0’s ability to generate condition-specific data opens new avenues for targeted in silico experiments and simulations.

MGM 2.0 also addresses critical challenges in personalized medicine, particularly in donor selection for FMT. By framing this problem as a question-answering task, the model effectively integrates recipient and donor information to predict post-transplant outcomes. The high ROUGE-1 scores achieved across datasets highlight the robustness of this approach, while in silico experiments demonstrate its potential for identifying “super donors” and optimizing transplant success. This innovation supports more precise donor selection, improving patient outcomes and minimizing risks associated with FMT.

Despite its strengths, one limitation of MGM 2.0 is that its effectiveness has been verified primarily in human microbiome problems, leaving its applicability to environmental microbiomes unexplored. While the contextual information derived from the sentence-like representation has proven sufficient for tasks involving human-associated microbiomes, future studies should evaluate the model’s performance in diverse environmental settings. Addressing this gap would help establish its broader utility across various ecological and industrial applications. Enhancing the Microbiome Turing Test with expanded metrics may also provide an even more rigorous evaluation of generative model performance.

In conclusion, MGM 2.0 represents a significant step forward in microbiome analysis harnessing the power of generative language models to transform how microbiome data is studied and applied. By bridging microbiome research and NLP, this sentence-like paradigm introduces new opportunities for data generation, predictive modeling, and precision medicine. The innovative methodologies and applications introduced by MGM 2.0 lay the groundwork for deeper insights, more robust predictions, and groundbreaking advancements in microbiome science.

## Methods

### Datasets and Preprocessing

#### Colonization dataset

Sequence data were retrieved from the European Nucleotide Archive (ENA) under study accession number PRJEB60398. Reads were preprocessed using fastp [34] according to the original study’s protocol: (1) Reads with more than 50% of bases below quality score 19 were removed. (2) Reads containing more than 5% N bases were removed. (3) Paired-end reads were discarded if read failed to meet the above criteria. Microbial community composition was then generated using metaphlan4 [35].

#### GMrepo

Abundance tables were downloaded from the GMrepo homepage (https://gmrepo.humangut.info/home). Diseases with fewer than 100 samples, along with samples labeled as “Infant,” “Premature,” and “Pregnant,” were excluded from the analysis.

#### FMT datasets

Abundance tables for the training set were obtained from the original study’s Zotero repository (https://doi.org/10.5281/zenodo.6611040). Sequence data for the two external datasets were retrieved from the Sequence Read Archive (SRA) under accession numbers PRJEB19232 and PRJNA380944. Microbial community composition for these datasets was generated using Qiime2 [3].

### Data Encoding

In this study, we employed a sentence-like approach to represent microbial community samples, transforming them into sequence-based representations for subsequent analysis. Each microbial community sample, denoted as {*x*_1_, *x*_2_, *x*_3_, …, *x*_*n*_} consists of a set of microbial taxa, where *x*_*i*_ represents the relative abundance of a specific genus in the sample. The vocabulary size, denoted as *X*, refers to the total number of unique genera observed across all samples in the dataset.

Next, the relative abundances *x*_*i*_ for each genus are normalized based on the mean and standard deviation of the MicroCorpus-260K dataset [15]. Let *μ*_*j*_ and *σ*_*j*_ represent the mean and standard deviation of the relative abundances of genus *j* across all samples in the dataset. The normalized relative abundance for genus *i* in a sample is calculated as:

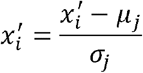

Following normalization, the genera in each sample are ranked based on their normalized relative abundance. The genus with the highest normalized abundance is assigned rank 1, and ranks decrease in descending order of abundance. The rank *r*_*i*_ of genus ii is defined as:

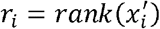

Finally, the ranks are tokenized into discrete input representations. Each rank *r_i_* is mapped to a unique token *t_i_*from a predefined vocabulary *V* ={*t*_*1*_ , *t*_*2*_ , …, *t*_*x*_ }, where *X* is the total number of unique genera across the entire dataset. This tokenization process transforms the ordered ranks into a sequence format, such that the microbial community sample is represented as a sequence of tokens:

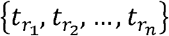

In this sequence, each token 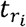 corresponds to the rank of genus *i*, capturing the relative structure of the microbial community while reducing reliance on the absolute abundance values.

## Model Architecture

We constructed MGM 2.0 model using eight layers of transformer blocks, with each block consisting of a self-attention layer and a feed forward neural network layer. Key hyperparameters were as follows: activation function, Gaussian Error Linear Unit (GELU); attention heads per layer, eight; embedding size: 256; feed forward size: 1024. The modeling framework was implemented in PyTorch, leveraging the Huggingface Transformers library for model configuration and training [36].

The self-attention mechanism employed in each transformer layer follows the scaled dot-product attention formula:

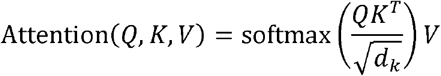

Where *Q* (queries), *K* (keys), and *V* (values) are the linear projections of the input, and *d*_*k*_ is the dimensionality of the keys. This formulation allows the model to capture contextual relationships between different tokens in the sequence, enabling more effective representation learning for downstream tasks.

## Downstream Fine-tuning

For downstream tasks, the pre-trained MGM model is fine-tuned by replacing the language modeling head with a task-specific head. All downstream tasks in this study focused on microbial community classification employed a sequence classification head, which utilized the final token (‘eos’ (end of sentence) in this study) for classification. Fine-tuning was executed using Huggingface’s Trainer API. Key hyperparameters included, learning rate, 1e-3, batch size, 50, warmup steps, 1000, weight decay, 0.001, validation split, 10% of the data.

## Prompt-guided Generation Process

To adapt the model for prompt-guided generation, we expanded the model’s vocabulary by introducing a new token immediately following the ’<bos>’ (beginning-of-sequence) token, designated as the label token. Let the extended vocabulary be denoted as *V*^′^ ={*t*_0_, *t*_1,_*t*_2, … ,_ *t*_*X* ,_*t*_*label*_}, where *t*_0_ corresponds to the ^’^<bos>^’^ token, *t*_1,_ *t*_2, …,_ *t*_*X*_ are the tokens representing the ranks of genera, and *t*_*label*_is the new token added to indicate the sample identity. Each sample in the microbiome dataset is then associated with a unique label token 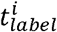, where *i* corresponds to the specific microbiome sample. For each microbiome sample *S*_*i*_={*x*_1_,*x*_2_,*x*_3_,…,*x*_n_}, we assign it a label token 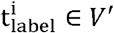 to represent the sample identity, so that the model can generate a sequence specific to that sample.

This vocabulary expansion ensures that the model can not only generate microbiome sequences but also tailor the sequences to reflect the characteristics of a specific sample. Formally, the input to the model is now a sequence of tokens 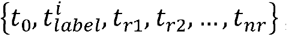, where 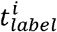 denotes the label token for the sample *i* and *t*_*r*1_,*t* _*r*2_,…,*t* _*rn*_ are the tokens representing the ranks of microbial genera in the sample.

Once this vocabulary expansion is made, the model is fine-tuned using next-token prediction. The training objective is to predict the next token in the sequence given the previous tokens. Let the current sequence of tokens be *t*_1_,*t*_2_ ,…,*t*_*k*_ , and the model’s goal is to predict the next token*t*_*k*+1_. The loss function used for training is the negative log-likelihood of predicting the orrect token *t*_*k*+1_, given the sequence of previously generated tokens:

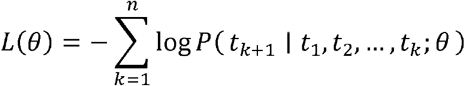

Here, *θ* represents the model parameters, and *P(t*_*k*+1_*t*_1_, *t*_2_,…, *t*_*k*_;)is the probability predicted by the model for the next token *t*_*k*+1_ conditioned on the previous tokens. This is learned by minimizing the cross-entropy loss over the entire sequence. During the generation phase, the model is provided with the label token *t*_*k*+1_ as the prompt, which signals the model to generate a sequence corresponding to the sample *i*. The model then generates a sequence of tokens 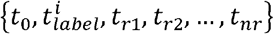 , where each *t*_*rk*_ represents the rank of the *k*^th^ genus in the sample

## Reconstructor Architecture

Once the model generates the sentence-like sequence, it is necessary to convert these sequences back into relative abundance values. To achieve this, we utilized the reconstructor network that was trained in parallel with the main model. The reconstructor’s task is to take the generated rank sequence and map it back to the original relative abundance scale. This step ensures that the final generated microbiome sequences are not only in the correct rank order but also reflect the original quantitative properties of the microbiome samples.

The reconstructor network was trained using a dataset of paired rank-encoded and abundance-encoded samples, learning to approximate the mapping from sentence-like representation back to relative abundance values. By using this approach, we preserve the diversity and abundance distributions of the microbiome, enabling the generation of realistic, contextually accurate microbiome data.

The reconstructor is a deep-learning model used to reconstruct microbial abundance. The ranked corpus after tokenizing was firstly encoded to a vector *X* ϵ [0,2]^*N*^,where

*X*_i_ =0 if species *i* is absent from this sample and *X*_*i*_ = *PE*(*i*) + 1 if it is present. Here, *PE*(*i*)is the position embedding from the Transformer with *d*_model_ = 1. The microbial composition of this sample is represented by a vector *y* ϵ [0,1) ^*N*^ , where *Y*_*i*_ is the relative abundance of species *i* in this sample. This deep-learning model (ReconstructorNet) was trained to learn the map from *x* to *y*. This model is a three-layer neural network with layers size *N* × 2*N* × *N* × *N*, using ReLU activation in the first two layers and Softmax in the final layer, optimized using Adam. The last two layers also employ residual connections to improve training stability and performance.

Pytorch-Lightning was used to build and train the model. Key hyperparameters included: Learning rate: 2e-4, Batch size: 64, Validation split: 20% of the data, Early stopping based on validation loss with 10 patience.

## Microbiome Turing Test

In the Microbiome Turing Test framework, we computed five first-order metrics and five second-order metrics. For the first-order metrics, cosine similarity, mean absolute error (MAE), Spearman correlation coefficient, alpha diversity and sparsity were calculated by comparing the generated data with the real data for each disease individually, and then averaging the results across 17 diseases to obtain the evaluation metric. Specifically, cosine similarity was computed using the cosine_similarity function from the sklearn.metrics.pairwise module, while MAE was calculated by taking the mean absolute differences between the real and generated data for each sample. Spearman correlation coefficients were determined using the spearmanr function from the scipy.stats module. Alpha diversity was assessed using Shannon entropy and Simpson index, calculated with the alpha_diversity function from the skbio.diversity module. Sparsity was evaluated using entropy, and the distributions of these entropy values were compared using the Wilcoxon signed-rank test, implemented in the scipy.stats module.

The second-order metrics focused on the biological significance of the data. For ROC-AUC, a Random Forest classifier trained on the real data was used to test the generated data from each model. The overlap of biomarkers was assessed by identifying the top 500 most important features in Random Forest classifiers trained on each dataset and then calculating the number of overlapping biomarkers between the real and generated data. Finally, we constructed Pearson correlation networks from the real and generated data using the networkx library. We evaluated network properties included network density, average degree, and modularity to determine how well the generated data captured the relationships between microbial communities. These processes were implemented using libraries such as numpy, pandas, torch, scipy, sklearn, and networkx.

## FMT Trial’s Representation

In this setup, we used a sequence-to-sequence (seq2seq) approach to model the transformation from the donor-recipient pair to the post-transplantation microbiome composition. Specifically, the input query consisted of the concatenated sentence-like representations of the recipient and donor microbiomes, with the two sets of representations separated by a special ‘sep’ token. The sentence-like representation of the post-transplantation microbiome sample was treated as the target sequence. Mathematically, the input for an FMT triad involving recipient *R*, donor *D*, and post-transplantation sample is *P* represented as:

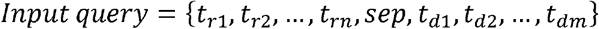

Where *t*_*r*1_,*t*_*r*2_,…,*t*_*rn*_ and *t*_*d*1_,*t*_*d*2_,…,*t*_*dm*_ represent the ranked tokens corresponding to the genera in the recipient and donor microbiomes. The ′ sep ′ token separates the recipient and donor representations. The target sequence is the sentence-like representation of the post-transplantation microbiome:

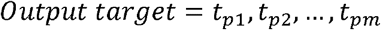

## Code Availability

The code for MGM model is available at https://github.com/HUST-NingKang-Lab/MGM.

## Acknowledgments

This work was partially supported by the National Key R&D Program of China (Grant No. 2023YFA1800900 and 2018YFC0910502), the National Natural Science Foundation of China (Grant Nos. 32071465, 31871334, 81827901). Numerical computations were performed on the Hefei Advanced Computing Center.

## Author contributions

KN conceived and proposed the idea. HZ and ZK designed and developed the framework. HZ, ZK, and YZ conducted the experiments, analyzed the data, and created the visualizations. RY provided valuable computing resources. HZ, ZK, YZ, RY, and KN contributed to editing and proofreading the manuscript. All authors reviewed and approved the final version of the manuscript.

## Competing interest

The authors declare that they have no competing interests.

## Ethics approval and consent to participate

Not applicable.

## Notes

### Competing Interest Statement

The authors have declared no competing interest.

