## Supplementary Materials for "Towards a Generative Paradigm for Large-scale Microbiome Analysis by Generative Language Model"

| Input Data | Research Question | Solution Approach |
| --- | --- | --- |
| Taxonomic profile abundance table (e.g., recipient microbiomes) | **Colonization Prediction**: Will a given exogenous species colonize the microbial community in a specific sample? | **Rank-based Representation**: Microbial taxa are ranked based on relative abundance, preserving structural relationships in the community. The model predicts colonization likelihood by leveraging the sentence classification model. |
| Taxonomic profile abundance table, disease labels (e.g., IBD, CDI) | **Data Simulation**: How can we generate realistic microbiome profiles conditioned on specific conditions, such as diseases? | **Prompt-Guided Data Generation**: The model generates synthetic microbiome data conditioned on disease-specific prompts (e.g., IBD or CDI labels). Fine-tuned generative models create realistic microbial profiles that reflect disease-associated microbial shifts. |
| Taxonomic profile abundance table (recipient and donor microbiomes) | **FMT Donor Selection**: Which donor microbiome is most suitable for transplant to a specific recipient to optimize health outcomes? | **Question-Answering (QA) Process**: The rank-based representations of both recipient and donor microbiomes are concatenated, with the model predicting the post-transplantation community composition. This allows predicting post-transplant microbiome composition from a recipient-donor pair and selecting the best donor based on predicted engraftment success (C2R values). |

**Supplementary Table 1: Overview of Methodological Approaches for Rank-based Representation Microbiome Tasks**

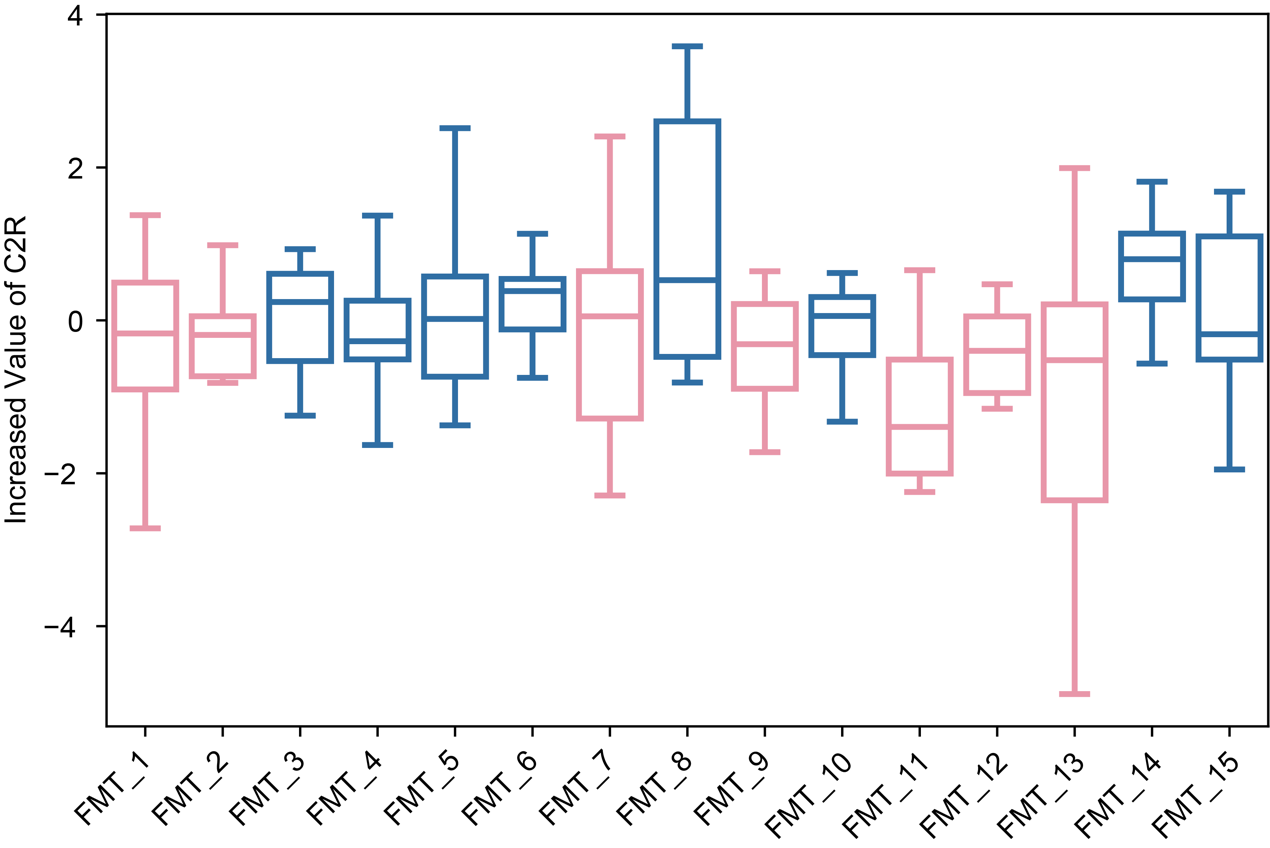

**Supplementary Figure 1. The increased value of C2R when assigned the donor to different recipients.** Pink value indicated the average C2R of the trials is below 0 while the blue value indicated the average C2R of the trials is above 0.

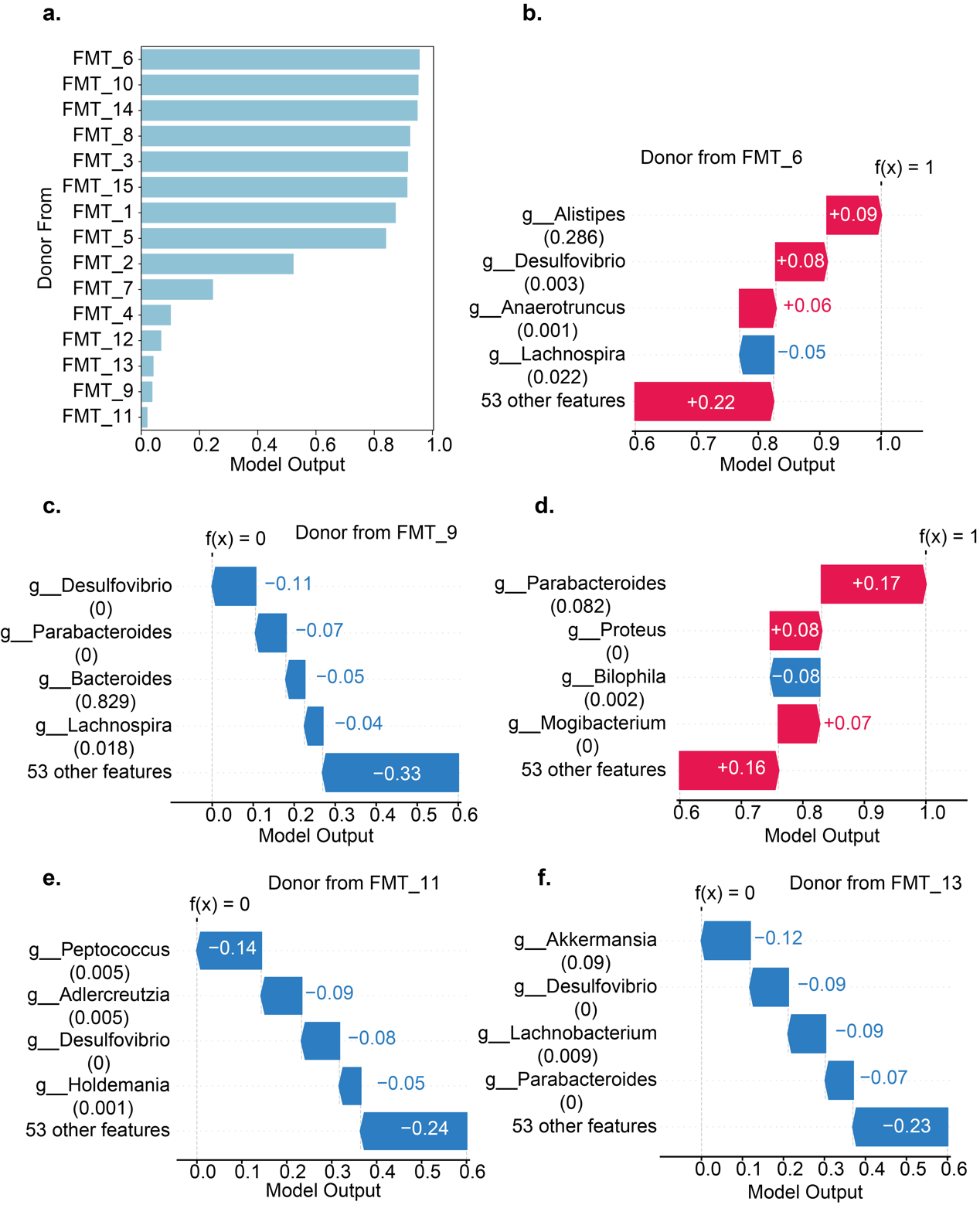

**Supplementary Figure 2. Results of SHAP analysis on donors from different FMT group. a.** The output of a LDA model to predict donors has positive or negative effect to recipient. **b.** Waterfall plot explaining the SHAP values for the donor from FMT14. **c.** Waterfall plot explaining the SHAP values for the donor from FMT14. **d.** Waterfall plot explaining the SHAP values for the donor from FMT14. **e.** Waterfall plot explaining the SHAP values for the donor from FMT14. **f.** Waterfall plot explaining the SHAP values for the donor from FMT14.
